# Evidence for distinct genetic and environmental influences on fear acquisition and extinction

**DOI:** 10.1101/2020.12.18.423504

**Authors:** K. L. Purves, G. Krebs, T. McGregor, E. Constantinou, K.J. Lester, T.J. Barry, M.G. Craske, K. S. Young, G. Breen, T. C. Eley

## Abstract

**Background:** Anxiety disorders are highly prevalent with an early age of onset. Understanding the aetiology of disorder emergence and recovery is important for establishing preventative measures and optimising treatment. Experimental approaches can serve as a useful model for disorder and recovery relevant processes. One such model is fear conditioning. We conducted a remote fear conditioning paradigm in monozygotic and dizygotic twins to determine the degree and extent of overlap between genetic and environmental influences on fear acquisition and extinction.

**Methods:** 1937 twins aged 22-25 years, including 538 complete pairs from the Twins Early Development Study (TEDS) took part in a fear conditioning experiment delivered remotely via the Fear Learning and Anxiety Response (FLARe) smartphone app.

In the fear acquisition phase participants were exposed to two neutral shape stimuli, one of which was repeatedly paired with a loud aversive noise, while the other was never paired with anything aversive. In the extinction phase the shapes were repeatedly presented again, this time without the aversive noise. Outcomes were participant ratings of how much they expected the aversive noise to occur when they saw either shape, throughout each phase.

**Results:** Twin analyses indicated a significant contribution of genetic effects to the initial acquisition and consolidation of fear, and the extinction of fear (15%, 30% and 15% respectively) with the remainder of variance due to the non-shared environment. Multivariate analyses revealed that the development of fear and fear extinction show moderate genetic overlap (genetic correlations .4-.5).

**Conclusions:** Fear acquisition and extinction are heritable, and share some, but not all of the same genetic influences.

## Introduction

Anxiety disorders affect over 20% of individuals during their lifetime (Kessler et al., 2012), have an early age of onset(Kessler et al., 2005), and are currently increasing sharply in prevalence (Baker, 2020; O’Connor, Downs, Shetty, & McNicholas, 2020). Both genetic and environmental influences are implicated in the development of anxiety, with twin heritability estimates ranging from 20-60% (Ask, Torgersen, Seglem, & Waaktaar, 2014; Meier et al., 2019; Polderman et al., 2015; Purves, Coleman, et al., 2019). Effective, evidence-based treatments exist for anxiety disorders, notably pharmacological approaches such as antidepressants and benzodiazepines (Baldwin et al., 2014), and psychological approaches such as Cognitive Behavioural Therapy (CBT) (Carpenter et al., 2018; Hofmann & Smits, 2008). However, only ∼50% of individuals respond regardless of treatment type (Clark et al., 2009; Cuijpers, Cristea, Karyotaki, Reijnders, & Huibers, 2016; Loerinc et al., 2015; Rush et al., 2006). Individual differences in treatment response are likely to be heritable, given known genetic influences on response to many aspects of the environment, such as life events and parenting (Kendler & Baker, 2007). Although there is evidence for familial clustering (Franchini, Serretti, Gasperini, & Smeraldi, 1998; O’Reilly, Bogue, & Singh, 1994) and influence of common genetic variants(Tansey et al., 2013) in response to antidepressant treatment for depression, the heritability of psychological treatment response is currently unknown. The largest studies in this field to date were insufficiently powered to draw strong conclusions (Coleman et al., 2016; Rayner et al., 2019).

Crucially, we do not yet know the specific mechanisms through which anxiety develops or recovery occurs, or how genetic influences contribute to these. Furthermore, using an experimental model of specific processes underlying disordered anxiety and treatment response has the advantage of a likely reduction in heterogeneity in the dataset (Scheveneels, Boddez, Vervliet, & Hermans, 2016). One possible set of experimental mechanisms is the acquisition and subsequent extinction of fears through direct learning experiences. These processes can be measured using fear conditioning paradigms. During the fear acquisition phase participants are exposed to two neutral stimuli, one of which is repeatedly paired with an aversive (e.g., a loud noise) or unconditional stimulus (US). The stimulus paired with the US is referred to as the CS+. The other stimulus is never paired with anything aversive and is referred to as the CS-. Over repeated trials, participants learn that the CS+ is associated with the aversive stimulus and typically demonstrate a ‘fear’ or ‘anxiety’ response to the CS+. This initial phase models processes implicated in anxiety development. Following acquisition, participants undergo an extinction phase where both CS+ and CS-are repeatedly presented without the aversive noise. The extinction phase models exposure-based treatment of fear and anxiety disorders used in CBT(Hofmann, 2008), which developed out of the fear extinction literature (Eelen & Vervliet, 2006).

A common way of measuring conditioning is to require participants to rate how much they expected the aversive outcome to occur with the CS+ and CS-throughout the task (Lonsdorf et al., 2017). These assessments can be thought of as risk-estimates. In order to measure task-related learning whilst controlling for general interindividual differences in responsivity, a differential metric (risk-estimates for the CS-subtracted from the risk-estimates for the CS+) is often calculated (Lonsdorf et al., 2017). In order to investigate the underlying contributions of genes and environment to these mechanisms, sample sizes are needed far in excess of those typically included in laboratory administered studies of fear conditioning. Notably, only one study in the largest meta-analysis of fear conditioning to date contained over 100 participants (Duits et al., 2015). One prior twin study obtained preliminary estimates of heritability of skin conductance responses during fear acquisition and extinction (0.20-0.46), and genetic overlap between them (Hettema, Annas, Neale, Kendler, & Fredrikson, 2003). However, although large in terms of fear conditioning, the sample size (173 twin pairs) was small for genetic analyses, as evidenced by large confidence intervals. Furthermore, the sample was split into two groups, each of which received a different stimuli set. Thus, whilst this provides preliminary evidence for the heritability of fear acquisition, extinction and their overlap, a larger, more heterogeneously administered study is required to consider this question.

The present study aimed to answer two important questions: (1) To what extent are fear acquisition and extinction influenced by genetic and environmental factors; (2) Are the genetic and environmental influences on the acquisition and extinction of fear shared between all processes, or are there distinct influences on each phase? We used the recently developed Fear Learning and Anxiety Response (FLARe) smartphone app (Purves, Constantinou, et al., 2019) to remotely administer a differential fear conditioning and extinction paradigm to a subset of twins from the Twins Early Development Study (TEDS) (Rimfeld et al., 2019). We assessed participant discrimination between the CS+ and CS-during early and late acquisition and extinction.

## Methods

### Sample

Participants were recruited by email invitation sent to 5934 twins enrolled in the Twins Early Development Study (Rimfeld et al., 2019), a longitudinal birth cohort study of twins born in England and Wales between 1994 and 1996. Twins who agreed downloaded the FLARe app (Purves, Constantinou, et al., 2019) via the iTunes or Google Play Stores. The experiment was completed by 2554 individuals of whom 1937 were included in the analyses after excluding those who removed their headphones, reduced phone volume<50%, or exited the app during the task. See **supplementary figure 1** for a detailed illustration of participant drop-out at each stage.

Twin zygosity was determined by parental responses to a twin similarity questionnaire. This has 95% concordance with genotyped zygosity(Price et al., 2000). The sample consisted of 250 complete monozygotic pairs (MZF=180, MZM=70), 288 dizygotic pairs (DZF=131, DZM=50, DZopp=107), and 860 singletons (F=549, M=312). See **supplementary table 1** for a description of the total sample including age and gender.

### Experimental Procedure

Participants were given instructions designed to maintain optimal experimental conditions, ensure headphone usage and maximum phone volume (see **supplementary figure 2-3**). They were asked to complete the task in a single, undisturbed session. The fear conditioning procedure began with a fear acquisition phase during which 12 presentations each of a large and small circle (conditional stimuli; CS) were shown in a pseudo-randomised sequence on a background image of an outdoor scene. One of these (CS+) was paired with the aversive ‘unconditional stimulus’ (US; a loud scream sound) during the final 500ms of the trial on 9 out of 12 presentations (75% reinforcement). The other was never paired with the US (CS-). The use of the large vs small circle as the CS+ was counterbalanced across participants. See **online supplement** for trial presentation rules. Participants then had a >10minute break, followed by a fear extinction phase. The same large and small circles were shown 18-times each on the background of an indoor living room scene. Neither shape was paired with the aversive stimulus. We used 18 trials in the extinction phase to ensure full extinction prior to the end of the task, given the remote delivery. After the extinction phase, participants were redirected to an external website(Qualtrics, 2019) to answer questions including whether or not they removed their headphones during the task. See **Figure 1** for a schematic of the experimental procedure.

**Figure 1.**
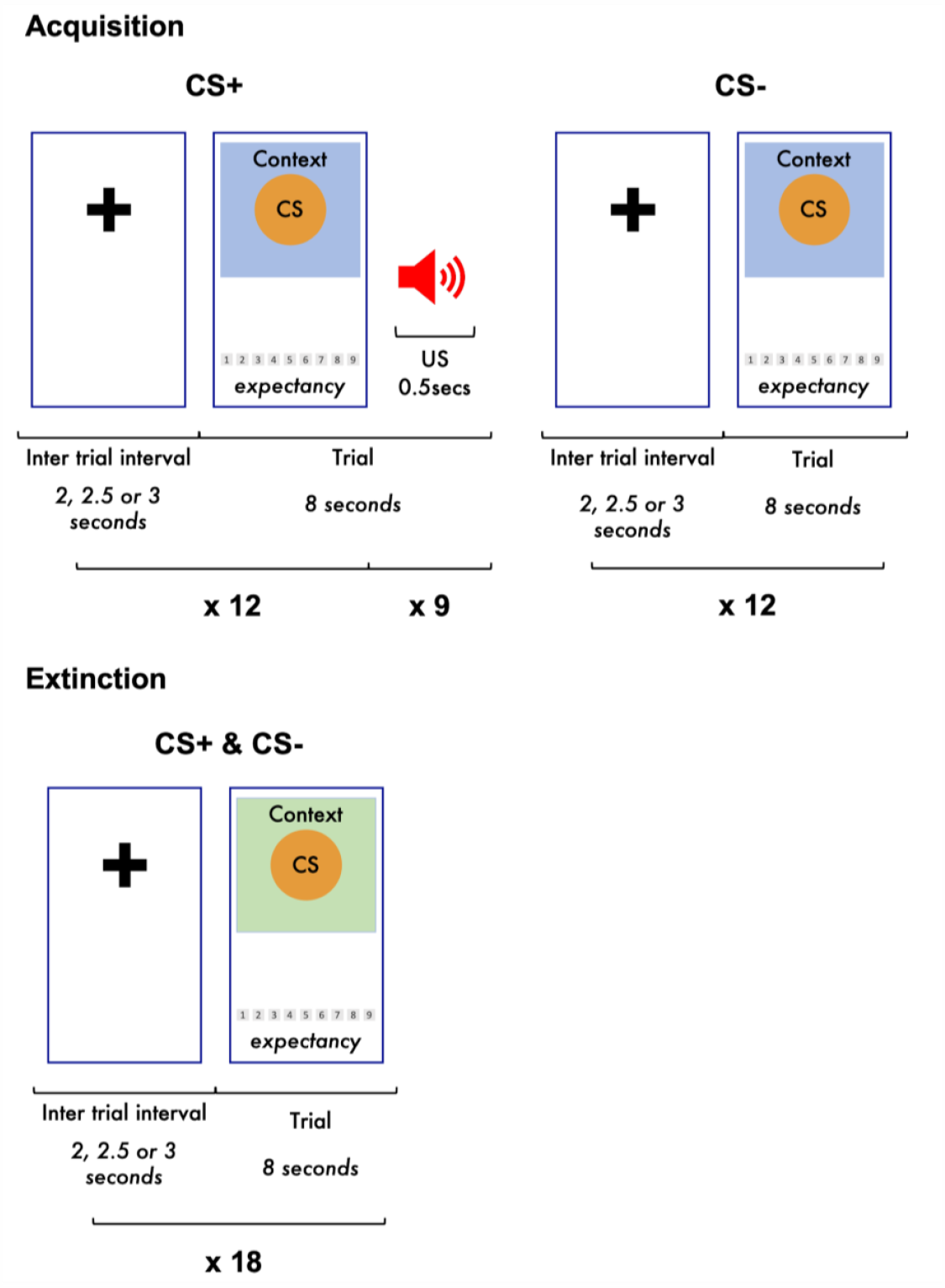
Schematic of fear conditioning procedures as implemented in the FLARe app Figure showing trial, and overall task structure of the fear conditioning task as implemented in the Fear Learning and Anxiety Response (FLARe) app. CS: conditional stimuli. Context: background image of an outdoor garden scene (acquisition) or indoor living room scene (extinction) displayed behind conditional stimuli during each trial. US: unconditional stimulus, a loud human scream played at maximum phone volume.

### Outcome Measures

During each trial, participants were asked to rate how much they expected to hear the aversive stimulus on a scale ranging from 1 (completely certain NO scream will occur) to 9 (completely certain a scream WILL occur). Risk-estimates (sometimes known as expectancy ratings) for the CS-were subtracted from those of the CS+ to obtain differential scores, an index of how well participants were able to differentiate between the two experimental cues. Repetition is a core component of memory and learning (Hintzman, 1976), and evidence from neuroimaging paradigms demonstrates that the association between fear conditioning and brain regions varies across time within phase (LaBar, Gatenby, Gore, LeDoux, & Phelps, 1998). Thus, in order to assess rapid acquisition of fear learning relative to later, consolidated fear learning after several repetitions, the post hoc decision was taken to consider fear acquisition and extinction in early and late stages. Differential scores created from risk-estimates made during the first and final thirds of the acquisition phase and the first third of the extinction phase were retained to assess the initial development, consolidation and extinction of fear learning respectively (see **Figure 2**). Differential scores for the last third of the extinction trials (late extinction) were not analysed in twin modelling as by this stage nearly all participants had reduced their risk-estimates for both stimuli to ∼1 (see **Table 1**).

**Figure 2.**
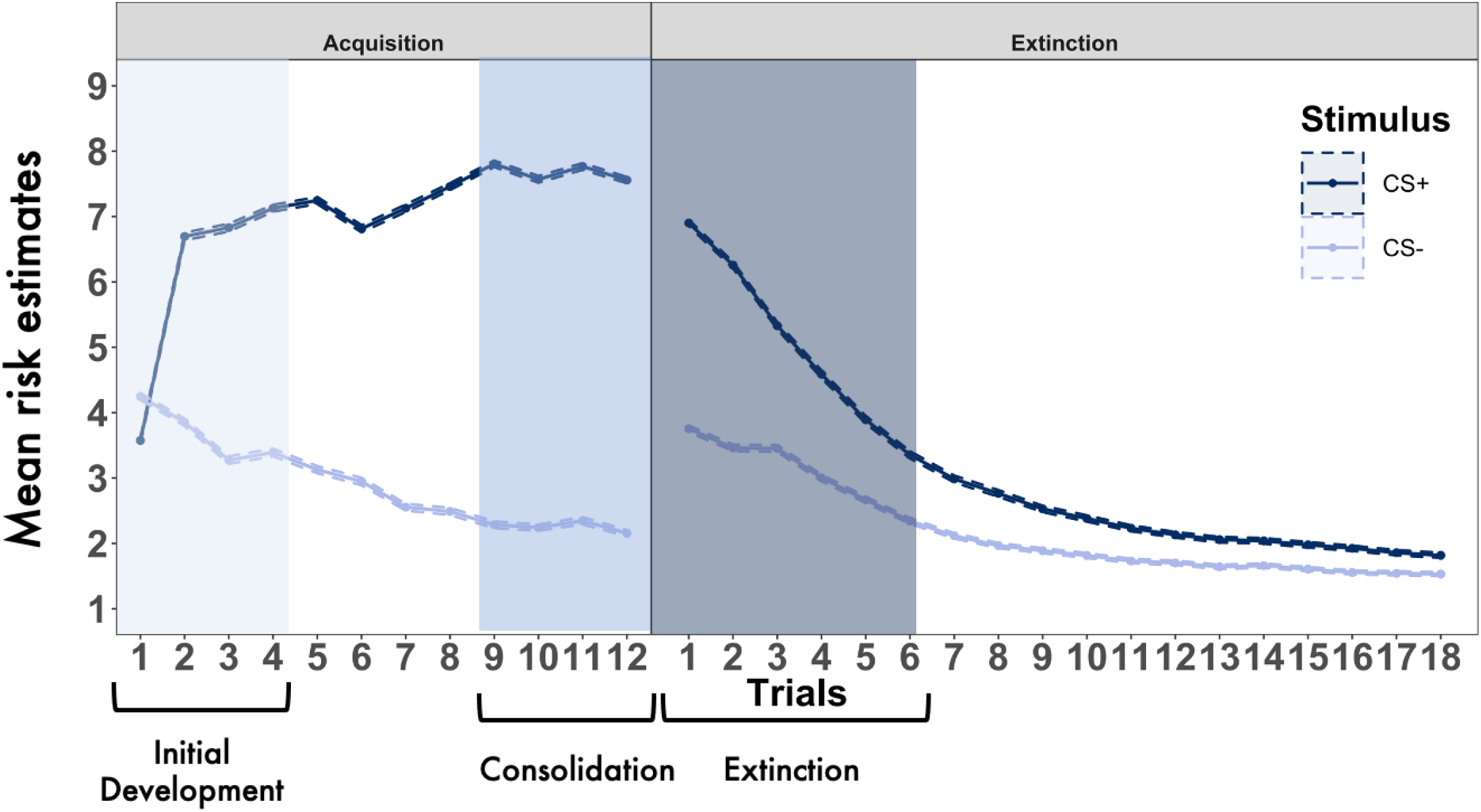
Average risk-estimates per trial across all participants during fear conditioning task Average risk-estimate per stimulus, per trial, averaged across all participants. Dashed shaded lines indicate standard error of the mean. CS+, conditional stimulus paired with the aversive scream; CS-, conditional stimulus never paired with the aversive scream.

**Table 1.** Descriptive statistics and twin pair intraclass correlations for fear conditioning outcomes

|  | <sup>a</sup>Average Risk Estimates |  |  | <sup>b</sup>Twin pair<br>Intraclass correlations for CS<br>differential |  | <sup>c</sup>CS+ / CS-<br>comparison |  |  |
| --- | --- | --- | --- | --- | --- | --- | --- | --- |
|  | CS+<br>mean (*SD*) | CS-<br>mean (*SD*) | CS differential<br>mean (*SD*) | MZ | DZ | t | df | p |
| Initial Development | 6.05 (*1.63*) | 3.70 (*1.56*) | 2.35 (*2.39*) | 0.14 | 0.08 | 34.26 | 1936 | < 2.2x10-16 |
| Consolidation | 7.67 (*1.74*) | 2.28 (*1.79*) | 5.39 (*3.16*) | 0.24 | 0.13 | 70.53 | 1936 | < 2.2x10-16 |
| Extinction | 5.06 (*1.85*) | 3.11 (*1.69*) | 1.95 (*2.12*) | 0.17 | 0.03 | 25.28 | 1936 | < 2.2x10-16 |
| Late Extinction | 1.96 (*1.68*) | 1.6 (*1.3*) | 0.36 (*1.23*) | - | - | 6.01 | 1936 | 2.2x10-09 |

Table comparing <sup>a</sup>mean and standard deviation (SD) for the average risk-estimate per stimulus (CS+ or CS-) during the first third of acquisition trials (Initial development), last third of acquisition trials (consolidation), first third of extinction trials (extinction) and the last third of extinction trials (late extinction), with <sup>b</sup>intraclass correlations between members of twin pairs for the <sup>c</sup>average difference in risk-estimates between the CS+ and CS- for each experimental phase (Initial development, Consolidation and Extinction). MZ; monozygotic twins DZ; dizygotic twins CS+; the conditional stimulus paired with the aversive scream CS-; the conditional stimulus that is never paired with the aversive scream CS differential; difference between CS+ and CS-

### Statistical analyses

#### Data preparation

Outcome variables (initial development, consolidation and fear extinction) were age and sex regressed to avoid artificial inflation of twin correlations (McGue & Bouchard, 1984), and the residuals were normalised using square-root transformation in R.

#### Multivariate twin modelling

Monozygotic (MZ) twins share 100% of their segregating genes, dizygotic (DZ) twins share ∼50%, while both MZ and DZ twins share their rearing environments (Rijsdijk & Sham, 2002). Monozygotic and dizygotic twin similarity can be compared to estimate genetic and environmental influences on a trait. The degree to which MZ twins are more similar than DZ twins reflects approximately half (100%-50%) of the genetic influence (labelled A; additive genetic influence). Twin resemblance not due to genetic factors is attributed to environmental factors that make twins more similar (common environmental factors, labelled C). Finally, the extent to which the MZ twin correlation differs from 1 reflects the non-shared environment and residual/error variance (labelled E).

A trivariate *Correlated Factors Solution* of the Cholesky Decomposition model was applied (see **figure 3**). This estimates the relative influence of latent factors A, C and E on each of the fear conditioning variables as well as the correlations between the genetic and environmental influences on each (Loehlin, 1996).

**Figure 3.**
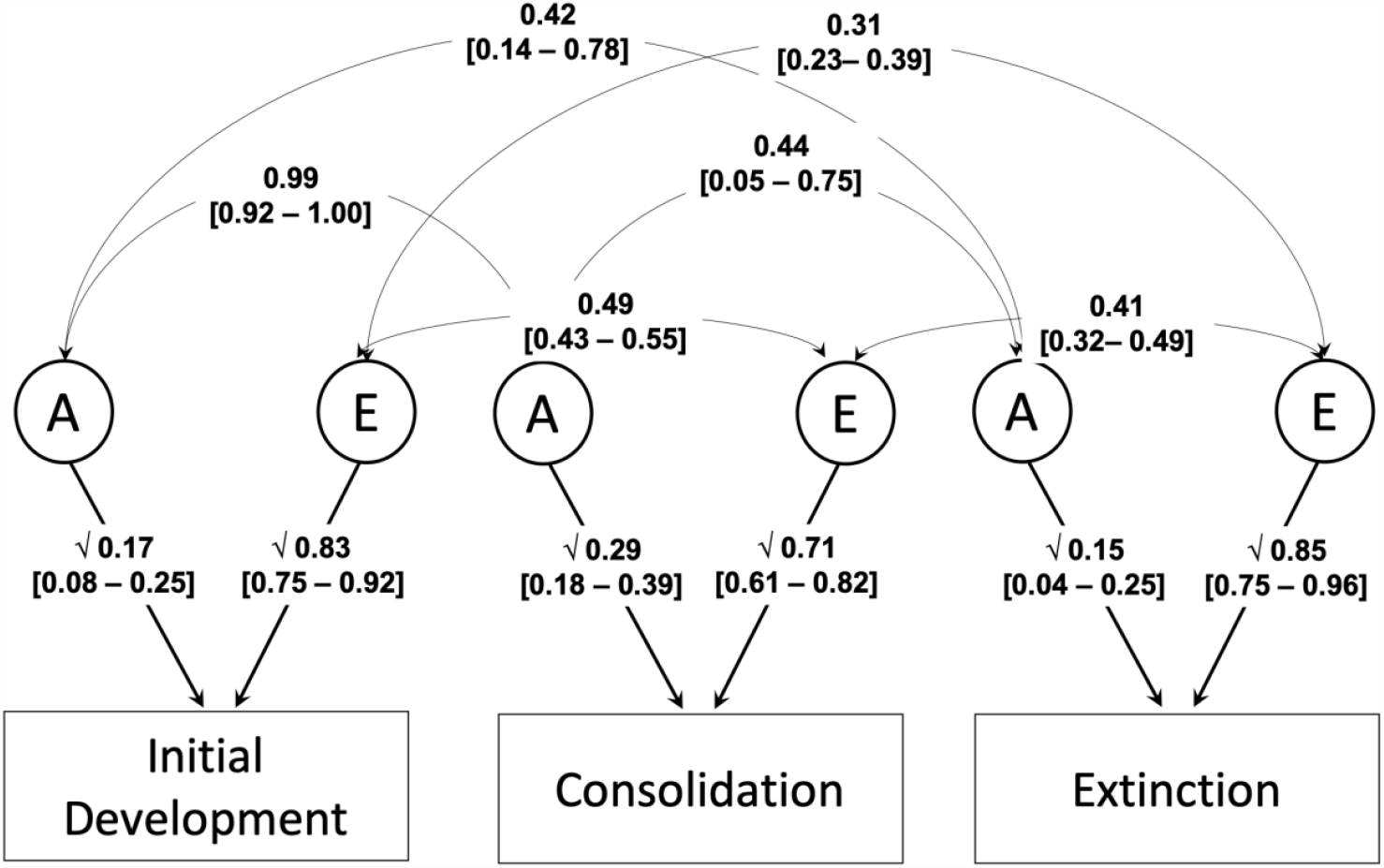
Trivariate correlated factors solution showing genetic and environmental influences on the initial development, consolidation and extinction of fear conditioning Figure shows the standardised path estimates and 95% confidence intervals for the AE trivariate correlated factors solution of the Cholesky model. A, Additive genetic effects; E, non-shared environment effects. Note that A and E present the proportion of phenotypic variance in each outcome accounted for by additive genetic and non-shared environment effects respectively. A and E for each outcome will sum to 100%. The curved paths show the correlations between the A and E factors for each outcome.

These parameters are then used to estimate the degree to which phenotypic correlations are attributable to genetic and environmental influences.

Genetic modelling was undertaken in R using the OpenMx package (Boker et al., 2011). OpenMx used Full Information Maximum Likelihood to estimate structural equation parameters, an effective method to deal with missing data (Boker et al., 2016; Jelicić, Phelps, & Lerner, 2009). Model fit was assessed using *X*^2^and Akaike Information Criterion (AIC).

## Results

### Differential fear conditioning

As shown in **Figure 2**, the difference between the CS+ and CS-was largest during the last 4 trials of acquisition, the phase we refer to as consolidation. The difference was smallest during late fear extinction when reinforcement of either stimulus had ceased. During the first four trials of acquisition (initial development), participants already showed a greater expectation of the scream occuring to the CS+ than the CS-, see first row, first section of Table 1 (Mean_CS+_=6.05, SD_CS+_=1.63, Mean_CS-_=3.70, SD_CS-_=1.56). By the consolidation phase (second row), risk-estimates had further increased for the CS+ and decreased for the CS-. Average risk-estimates reduced for both stimuli over the course of extinction. During the first third of extinction trials, ratings were similar to average risk-estimates during the initial development of fear. By the end of extinction, almost all participants had ceased to report any expectation of a scream occurring for either stimulus.

### Trivariate twin modelling

See **Table 1** middle section for monozygotic and dizygotic twin correlations for each fear conditioning variable and **Table 2** for cross-twin cross-trait genetic and phenotypic correlations. We only report results from multivariate modelling, as these are better powered than univariate models. For five of the six pairs of correlations, the ACE model produced C estimates that were non-significant and close to zero, and we therefore tested a more parsimonious AE model, which did not result in a significant loss of fit and was therefore selected as the final model (see **supplementary table 2** for model fit information).

**Table 2.** Cross-twin cross-trait (lower) and phenotypic (upper) correlations with proportion of variance explained by A and E

|  | Initial Development | Consolidation | Extinction |
| --- | --- | --- | --- |
| Initial Development | **0.32**<br>*[0.08 – 0.25]* | **0.59**<br>*[0.55 – 0.64]*<br>A 37% \| 63% E | **0.32**<br>*[0.28 – 0.38]*<br>A 19% \| 81% E |
| Consolidation | **0.21**<br>*[0.13 – 0.29]*<br>**0.12**<br>*[0.05 – 0.20]* | **0.29**<br>*[0.18 – 0.39]* | **0.38**<br>*[0.34 – 0.43]*<br>A 22% \| 78% E |
| Extinction | **0.05**<br>*[-0.05 – 0.15]*<br>**0.04**<br>*[-0.05 – 0.15]* | **0.08**<br>*[0.00 – 0.17]*<br>**0.03**<br>*[-0.04 – 0.13]* | **0.15**<br>*[0.04 - 0.25]* |

Table showing cross-twin cross-trait correlations [95% confidence interval] for MZ (top) and DZ (bottom) twin pairs (below diagonal) and phenotypic correlations [95 % confidence interval] with the standardised proportion of phenotypic association accounted for by A / E (above diagonal). Estimate of heritability from multivariate twin analyses [95% confidence interval] shown on the diagonal. Confidence intervals are obtained by bootstrapping over 100 iterations.

Estimates from this model are shown in **Figure 3**. Initial development, consolidation and extinction of fear were all significantly influenced by additive genetic factors (A). For example, the estimate of heritability (a^2^) for consolidation was 0.29 (0.18-0.39). In addition, there were significant shared genetic effects (r_a_) between all variables, with genetic correlations being highest between initial development and consolidation of fear (rG=0.99 [0.97-1.00]). In comparison, genetic correlation were significantly lower between each of the acquisition phases and extinction of fear (initial development and extinction: rG=.42 [.14 - .78]; consolidation and extinction: rG = .44 [.05 - .75]). The effect of non-shared environment (E) was also significant for all variables (range e^2^=0.71 - 0.83), with significant non-shared environmental correlations (0.31-0.49) between all variables. See **Table 2** for the standardised proportion of the phenotypic variance between variables accounted for by A and E respectively.

## Discussion

We present the largest twin study of fear conditioning to date, providing evidence for genetic and non-shared environmental influences. Discriminative fear learning is significantly heritable during the initial development, consolidation and extinction of fear conditioning. We demonstrated significant contributions from non-shared environmental influences (which includes measurement error), but no evidence for the contribution of common environment. Shared genetic effects between initial development and the consolidation of fear approached unity. This indicates that the genes involved in the initial development of fear learning are virtually the same as those involved in the later consolidation of fear learning. The genetic correlations between both stages of fear acquisition and extinction were moderate and significant, indicating that some of the same genes implicated in fear learning also play a role in the extinction of these learned associations. However, these genetic correlations were around 0.5, indicating that roughly half the genetic factors on extinction of fear are different from those on fear learning. Whilst a previous study provided preliminary evidence for complete genetic overlap between acquisition and extinction of fear, these results were difficult to interpret as confidence intervals were not included (Hettema et al., 2003). Our study thus provides the first robust evidence for the role of genes in the acquisition and extinction of fear.

Findings should be considered in light of some limitations. First, the remote delivery of the experiment necessitates lower experimental control compared to laboratory delivery. This is particularly reflected in the number of individuals who removed their headphones during the task, and thus excluded from analyses. Second, the sample was drawn from a population based twin cohort. In future studies, it will be important to test whether there are differences in the genetic and environmental contributors to fear conditioning between individuals with anxiety disorders compared to healthy individuals.

### Genetic influences on fear acquisition

We explored genetic influences on two stages of fear acquisition. Initial fear development was calculated using risk-estimate differences for the CS+ and CS-during the first third of acquisition trials. Fear consolidation was indexed by the risk-estimates differences during the final third of trials. Although initial development and consolidation shared all of the same genetic influences, genetic factors accounted for more variance in fear consolidation. A persistent reduced capacity to distinguish between cues paired and never paired with threat is similar to the tendency to interpret neutral stimuli as threatening (negative interpretation bias). This cognitive bias is a core feature of anxiety in adults and children (Bar-Haim, Lamy, Pergamin, Bakermans-Kranenburg, & van IJzendoorn, 2007; Dudeney, Sharpe, & Hunt, 2015; Lau & Waters, 2017; Lester, Lisk, Carr, Patrick, & Eley, 2019). Generalised Anxiety Disorder (GAD) in particular is associated with interpretation biases (M W Eysenck, MacLeod, & Mathews, 1987; Michael W Eysenck, Mogg, May, Richards, & Mathews, 1991; Hayes, Hirsch, Krebs, & Mathews, 2010; Mathews, Richards, & Eysenck, 1989; Mogg et al., 1994), which we have previously shown to be heritable (Brown et al., 2014; Eley et al., 2008; Lau, Belli, Gregory, & Eley, 2014). In sum, the consolidation of fear learning is probably more relevant than initial fear learning to understanding anxiety disorders. Notably, the heritability of fear consolidation (29%) is similar to estimates of generalised anxiety disorder from previous twin studies (Hettema, Neale, & Kendler, 2001). This supports the possibility that the consolidation of fear conditioning is a relevant mechanism underlying disordered anxiety.

### Genetic and environmental influences on fear extinction

The modest but significant heritability of fear extinction (15%) suggests that a relatively small degree of variation in fear extinction in healthy individuals is influenced by genetic factors. There have been no twin studies of treatment response for anxiety, and two genome-wide meta analyses of Cognitive Behavioural Therapy (CBT) treatment response have failed to derive significant heritability estimates despite having statistical power to detect heritability of 30% or greater (Coleman et al., 2016; Rayner et al., 2019). This study thus adds to our limited knowledge about the relative role for genes and the environment in fear extinction, a mechanism underpinning exposure-based treatment, a core component of CBT for anxiety disorders.

The high estimate of the non-shared environment (e^2^ = 85%) influencing fear extinction in this study reflects both high measurement error given our task (Purves, Constantinou, et al., 2019), and true environmental influence. There are several plausible contributors to variable treatment response over and above direct genetic influences. These include specific life experiences, and individual characteristics such as unemployment, low educational attainment and poor interpersonal relationships that are also known to reduce treatment efficacy (DeRubeis et al., 2014; Mojtabai, 2017; Newman, Llera, Erickson, Przeworski, & Castonguay, 2013; Renaud, Russell, & Myhr, 2014). Specific disorder profiles, including greater symptom severity, comorbidity with other mental health disorders and poor treatment adherence are associated with poor treatment response in anxious adults and children (Hudson et al., 2013, 2015; Rayner et al., 2019; Wergeland et al., 2016). Understanding modifiable personal risk factors may help identify areas for intervention that could serve as a precursor or adjunct to therapy to further enhance the effect of exposure within a personalised medicine framework.

### Genetic and environmental overlap between fear acquisition and extinction

There were moderate genetic (r_a_=0.41-0.44) and non-shared environment (r_e_=0.31-0.41) correlations between fear acquisition and extinction. If fear extinction is considered a credible marker underlying treatment response (Craske, Hermans, & Vervliet, 2018),then at least some of the genetic markers underlying treatment response are the same as those underlying anxiety development, and approximately half the genes relevant to extinction should be identified in GWAS of anxiety. This “cause informs cure” (Uher, 2008) perspective would mean that findings from the rapidly growing anxiety genetics field would be relevant to understanding genetic influences on psychological treatment response.

We note genetic correlations between fear acquisition and extinction were below 0.5, indicating that distinct influences exist. Crucially, this indicates understanding the genetics of anxiety disorder is not sufficient to fully understand the genetics of psychological treatment response. It continues to be important to find ways of investigating treatment response as a distinct phenotype in large samples of individuals with sufficient depth of phenotyping to explore the role of variance in a range of environmental and individual factors within genetically informed designs.

## Conclusions

We presented robust evidence for genetic influences on fear acquisition and extinction, shedding light on possible mechanisms underlying known genetic influences on anxiety. We have also shown moderate genetic overlap between fear acquisition and extinction. This indicates that genetic susceptibility to developing fears is likely to have a moderate influence on reducing learned fears, for example through exposure-based treatment. Future studies using genotyped samples with fear conditioning or treatment outcome data will enable more detailed investigation of the genetics of the development and treatment of anxiety.

In conclusion, these findings represent a significant advance in the genetics of fear acquisition and extinction, two key processes underlying the development and exposure-based treatment of anxiety disorders.

## Ethical standards

The authors assert that all procedures contributing to this work comply with the ethical standards of the relevant national and institutional committees on human experimentation and with the Helsinki Declaration of 1975, as revised in 2008.

