## Supplemental material for "Evidence for distinct genetic and environmental influences on fear acquisition and extinction"

### Supplementary figures and tables

**Supplementary figure 1.** Number of participants invited and successfully completing the experiment

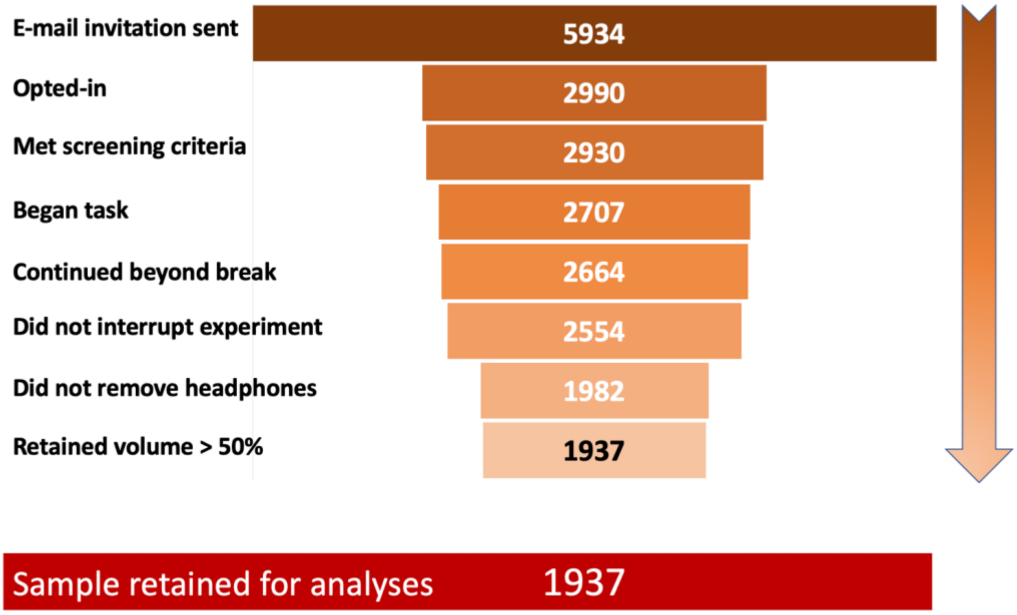


The figure shows the sample size from recruitment to final analyses. 'Opting-in' to the study involved downloading and completing informed consent procedures. 'Continued beyond break' indicates that participants returned after the ten-minute break between acquisition and extinction. 'Did not interrupt the experiment' indicates that the participants did not exit or close the app during the task. 'Did not remove headphones' indicates that they did not report removing their headphones during the task. 'Retained volume > 50%' refers to the phone volume during the task in the fear acquisition phase.

**Supplementary figure 2.** Pre-task and setup instructions

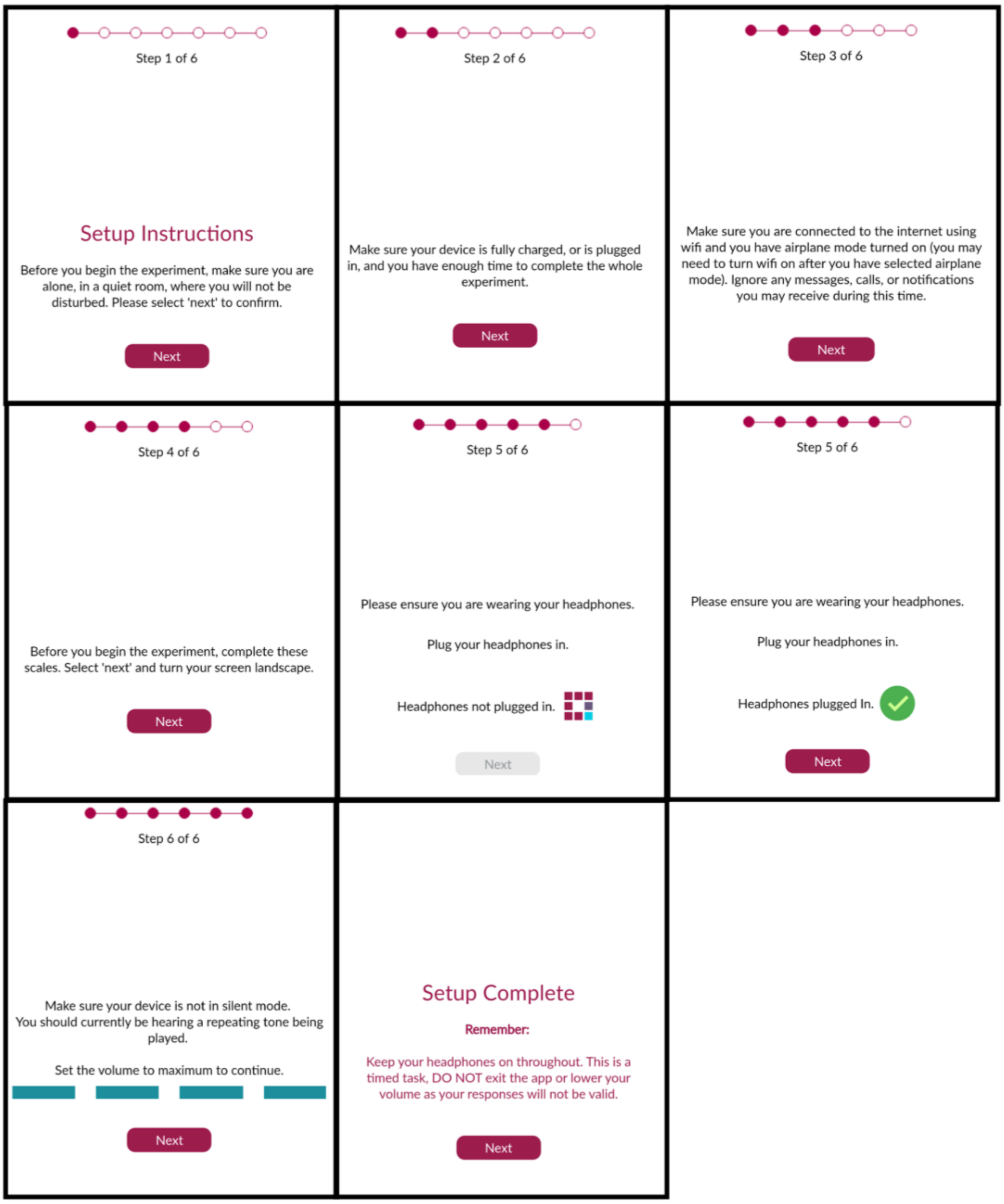


Screenshots of the initial pre-task set-up instructions for the Fear Learning and Anxiety Response (FLARe) app in the order which they are seen by participants.Participants navigate by clicking the buttons labelled 'next' at the bottom of the screen. Where a button is grey, participants are unable to click it to proceed to the next screen. This indicates that some action is required by the participant in order to progress. These screens are seen after consent and screening procedures, but before the task begins.

**Supplementary figure 3.** Fear conditioning task instructions

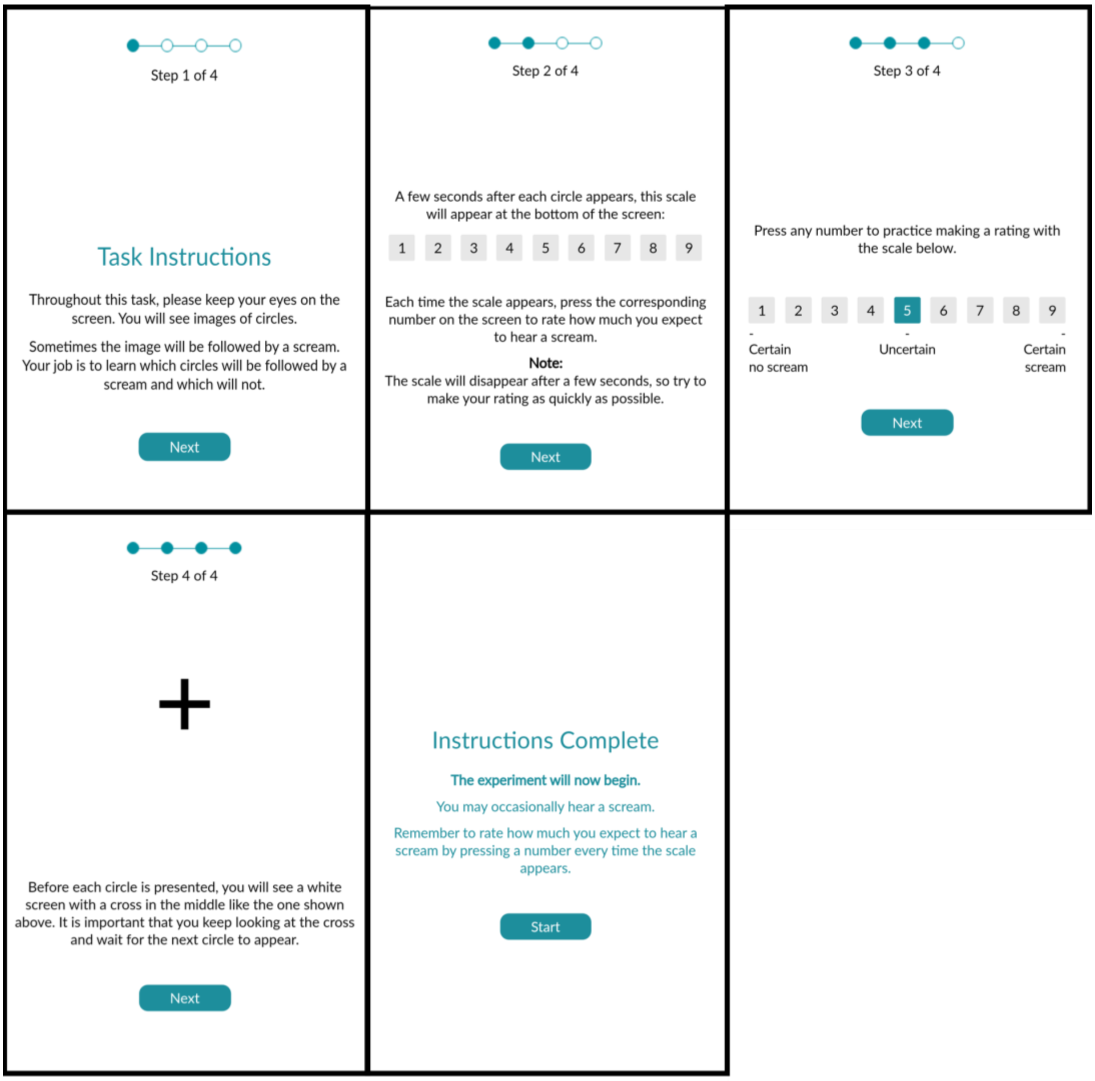


Screenshots of the task instructions as they appear during participation in the Fear Learning and Anxiety Response (FLARe) app. Screens are shown in the order which they appear. Participants navigate by clicking the buttons labelled 'next' at the bottom of the screen. Screens commenced immediately after the initial set-up was completed, and the fear conditioning task began after the final screen.

**Supplementary table 1.** Demographic and sample characteristics

| | Monozygotic twins | Dizygotic twins | Whole sample |
|---|---|---|---|
| **Total sample size (n complete pairs)** | 791 (250) | 1145 (288) | 1937 (538) |
| **n female (%)** | 559 (71%) | 766 (67%) | 1325 (68%) |
| **Mean age [range]** | 23.70 [22.20 - 25.17] | 23.51 [22.20 - 25.20] | 23.67 [22.20 - 25.17] |

Table shows the total number of complete twin pairs and singletons included in the study by zygosity. Mean age and proportion of the sample that are female are presented for each group. Note that none of the differences between monozygotic and dizygotic twin pairs were statistically different.

**Supplementary table 2.** Goodness of fit statistics for the trivariate twin analysis of all variables

| | | | Model fit | | | Difference in fit | | |
|---|---|---|---|---|---|---|---|---|
| **Model** | **Description** | **Sources of variance** | **$\chi^2$** | **df** | **AIC** | **$\Delta\chi^2$** | **$\Delta$df** | **p** |
| ACE | Full model | $A^1C^1E^1$ - $A^2C^2E^2$ – $A^3C^3E^3$ | 21194.26 | 5898 | 9398.26 | | | |
| AE | Constrain C to zero | $A^1E^1$ - $A^2E^2$ - $A^3E^3$ | 21194.26 | 5904 | 9386.26 | 0.004 | 6 | 1.00 |

Table showing the goodness of fit statistics for the trivariate twin analyses of all variables. A, Additive genetic effects; C, Shared environment effects; E, non-shared environment effects. $\chi$2, chi-squared likelihood ratio test; df, degrees of freedom; AIC, Akaike's Information Criterion; $\Delta\chi$2, difference in chi-squared likelihood ratio between models; $\Delta$df, difference in degrees of freedom between models; p, significance test for the $\Delta\chi$2. $p < 0.05$ would indicate a significantly worse fit of the model.

### Supplementary text

#### Trial presentation rules

During the fear conditioning acquisition trials, the following rules for trial order applied: The first two trials were a CS+ and CS- (order randomised across participants), the first CS+ was always paired with the aversive US and there were no more than two consecutive trials with the same CS. Between trials, an intertrial interval of a random duration of 2, 2.5 or 3 seconds occurred, during which a black cross was shown on a white screen. During extinction, The first two trials were a CS+ and CS-  (order randomised across participants) and there were no more than two consecutive trials with the same CS.
